# Metagenomic profiling of ticks: identification of novel rickettsial genomes and detection of tick borne canine parvovirus

**DOI:** 10.1101/407510

**Authors:** Anuradha Ravi, Suheir Ereqat, Amer Al-Jawabreh, Ziad Abdeen, Omar Abu Shamma, Holly Hall, Mark J. Pallen, Abedelmajeed Nasereddin

## Abstract

**Background:** Across the world, ticks act as vectors of human and animal pathogens. Ticks rely on bacterial endosymbionts, which often share close and complex evolutionary links with tick-borne pathogens. As the prevalence, diversity and virulence potential of tick-borne agents remain poorly understood, there is a pressing need for microbial surveillance of ticks as potential disease vectors.

**Methodology/Principal Findings:** We developed a two-stage protocol that includes 16S-amplicon screening of pooled samples of hard ticks collected from dogs, sheep and camels in Palestine, followed by shotgun metagenomics on individual ticks to detect and characterise tick-borne pathogens and endosymbionts. Two ticks isolated from sheep yielded an abundance of reads from the genus *Rickettsia*, which were assembled into draft genomes. One of the resulting genomes was highly similar to *Rickettsia massiliae* strain MTU5. Analysis of signature genes showed that the other represents the first genome sequence of the potential pathogen *Candidatus* Rickettsia barbariae. Ticks from a dog and a sheep yielded draft genome sequences of strains of the Coxiella-like endosymbiont *Candidatus* Coxeilla mudrowiae. A sheep tick yielded sequences from the sheep pathogen *Anaplasma ovis*, while *Hyalomma* ticks from camels yielded sequences belonging to *Francisella*-like endosymbionts. From the metagenome of a dog tick from Jericho, we generated a genome sequence of a canine parvovirus.

**Significance:** Here, we have shown how a cost-effective two-stage protocol can be used to detect and characterise tick-borne pathogens and endosymbionts. In recovering genome sequences from an unexpected pathogen (canine parvovirus) and a previously unsequenced pathogen (*Candidatus* Rickettsia barbariae), we demonstrate the open-ended nature of metagenomics. We also provide evidence that ticks can carry canine parvovirus, raising the possibility that ticks might contribute to the spread of this troublesome virus.

**Author Summary:** We have shown how DNA sequencing can be used to detect and characterise potentially pathogenic microorganisms carried by ticks. We surveyed hard ticks collected from domesticated animals across the West Bank territory of Palestine. All the ticks came from species that are also capable of feeding on humans. We detected several important pathogens, including two species of *Rickettsia*, the sheep pathogen *Anaplasma ovis* and canine parvovirus. These findings highlight the importance of hard ticks and the hazards they present for human and animal health in Palestine and the opportunities presented by high-throughput sequencing and bioinformatics analyses of DNA sequences in this setting.

## Introduction

Ticks are ectoparasitic arthropods that feed exclusively on the blood of their vertebrate hosts. Across the world, ticks act as vectors of human and animal pathogens (including viruses, bacteria and protozoa), often mediating transfer of infection from one host species to another, including zoonotic infections of humans [1–3]. Aside from negative effects on human and animal morbidity and mortality, ticks and tick-borne diseases are responsible for huge global production losses, amounting to US$ 14-19 billion per annum [4].

Many tick-borne infections remain undiagnosed and the prevalence, diversity and virulence potential of tick-borne agents remain poorly understood. Furthermore, as most tick-borne pathogens are hard to grow in the laboratory and the ticks themselves may be hard to identify on morphological grounds, our understanding of pathogen and tick population structure and the fine-grained genomic epidemiology of tick-borne infections remains patchy. These gaps in our knowledge highlight the need for microbial surveillance in tick vectors to assess the risk of infection in humans and animals.

Ticks rely on bacterial endosymbionts to provide nutrients lacking from their highly restricted blood-centred diet, as evidenced by the observation that ticks treated with antibiotics show decreased fitness at all life stages [5–7]. Interestingly, some intracellular endosymbionts of ticks share close and complex evolutionary links with tick-borne pathogens of humans, such as *Coxiella burnetii* and *Francisella tularensis,* with *Coxiella*-like and *Francisella*-like endosymbionts (CLEs and FLEs, respectively) occurring in a wide range of ticks worldwide [8–20].

Transitions between pathogenic and symbiotic lifestyles occur in both directions—thus, *Coxiella burnetii* is thought to have evolved from a vertically transmitted tick endosymbiont, whereas most FLEs probably evolved from pathogenic strains of *Francisella* [11,13,21]. Interactions with symbionts influence the abundance of pathogens within ticks. For example, *Francisella*-like endosymbionts inhibit colonisation with the spotted fever group rickettsias and with *Francisella novicida*, while the endosymbiont *Rickettsia bellii* is inversely associated in abundance with *Anaplasma marginale* [22]. Similarly, in North American ticks, the obligate intracellular endosymbiont *Rickettsia peacockii*, inhibits its pathogenic relative *R. rickettsia*, the causative agent of Rocky Mountain Spotted Fever [23].

Surveillance of ticks using taxon-specific PCR assays has shed light on the distribution of tick-borne pathogens and endosymbionts in Palestine and in the neighbouring state of Israel. We detected *Bartonella* in nearly 4% of hard ticks and spotted-fever-group *Rickettsia* in 17% of hard ticks collected from domesticated animals in Palestine [24,25]. Similarly, we have found apicomplexan parasites including *Theileria, Babesia*, and *Hepatozoon* in ixodid ticks from Palestine [26]. Various pathogens have been detected in ticks collected in Israel, including *Ehrlichia canis, Anaplasma spp., Babesia canis vogeli, Rickettsia massiliae, R. sibirica mongolitimonae, R. africae, R. aeschlimannii,* [27–31]. *Rickettsia massiliae* is widely disseminated throughout Israel in questing ticks, unattached to animals [32]. Endosymbionts have also been described in ticks from Israel, including the mitochondrial endosymbiont *Midichloria mitochondrii* [30]. *Francisella*-like endosymbionts have been reported in over 90% of *Hyalomma* ticks obtained from domesticated animals and migratory birds in Israel [12], while *Coxiella*-like endosymbionts have been identified and characterised in local *Rhipicephalus* ticks [33,34].

Although amplification and sequencing of taxon-specific PCR products has proven useful in microbial surveillance of ticks, this approach provides limited information on the target microorganisms, plus it only finds what has been targeted in the assay. Use of molecular barcodes such as 16S ribosomal gene sequences allows a more open-ended approach, capable of detecting numerous species in the same sample [24]. However, this approach provides limited insights into the biology, evolution and spread of the species and lineages under examination. The advent of high-throughput sequencing offers the potential for a powerful new open-ended metagenomics approach to the identification and characterisation of pathogens and endosymbionts [35]. This approach has already been used to identify novel viruses and to detect bacterial pathogens and symbionts in ticks [34,36]. It also brings the promise of sequence-based identification of ticks and hosts. However, widespread application of this approach is still limited by cost.

Here, we present a two-stage tick surveillance strategy that combines an initial round of 16S-amplicon screening of pooled tick samples, followed by focused shotgun metagenomics investigations, which confirmed the identities of the ticks as well as providing phylogenomic information on tick pathogens and endosymbionts, including the first genome sequence of the potential pathogen *Candidatus* Rickettsia barbariae.

## Materials and Methods

### Study design and sample collection

We employed a two-stage protocol that includes 16S-amplicon screening of pooled tick samples followed by focused shotgun metagenomics investigations on individual ticks (Fig. 1).

**Fig. 1.**
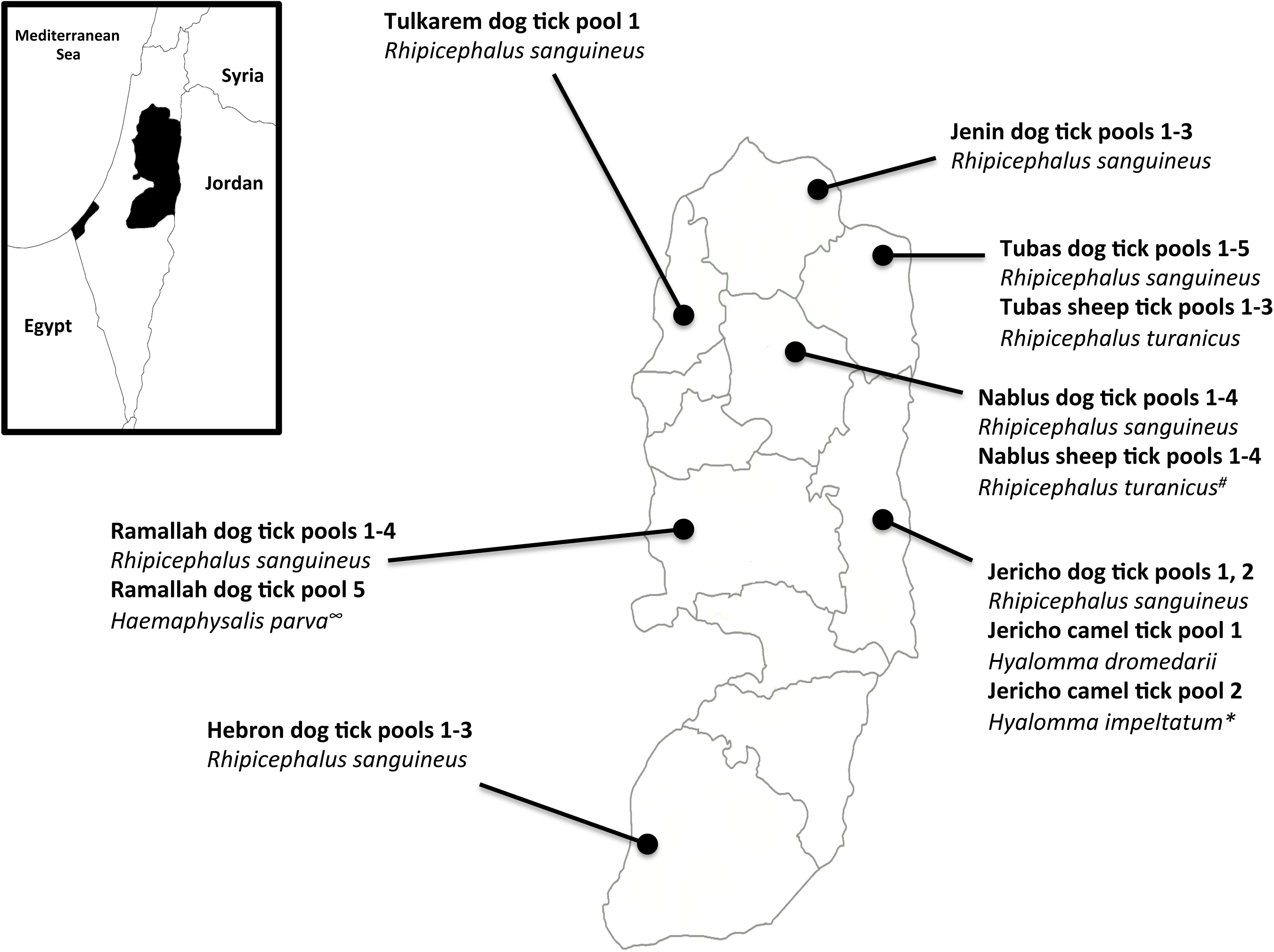
Study workflow.

Hard ticks were collected in 2015 from domesticated animals from seven governorates of the West Bank, Palestine, including: two camels from Jericho; four sheep from Nablus; three sheep from Tubas; and 23 dogs from all seven study sites (Fig. 2). Ticks were gently removed from host animals by forceps or hand and individually placed into small, labeled plastic tubes containing 70% ethanol for morphological identification. Ticks were identified using standard taxonomic keys and stored at −20°C until DNA extraction [24]. The animal owners were verbally informed about the study objectives and sampling procedure. All owners gave their verbal informed consent to collect ticks from their animals. The ethics committee at Al-Quds University approved the study (EC number: ZA/196/013).

**Figure 2.**
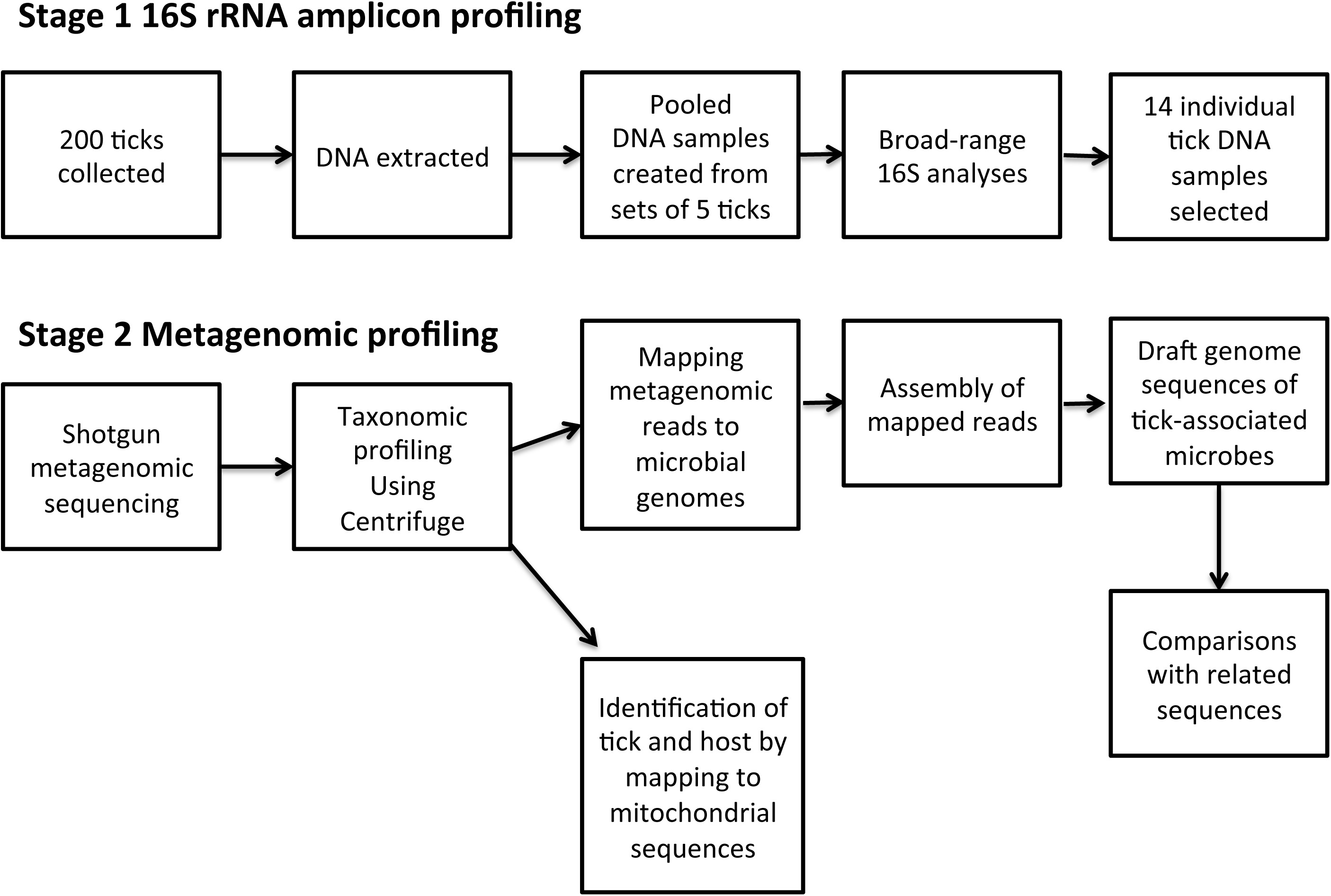
Collection sites of pooled tick specimens from the West Bank, Palestine. Species identities according to morphological criteria. \**Hyalomma dromedarii* according to sequence identity; ∞unidentified *Haemaphysalis* sp. according to sequence identity; # Nablus sheep tick 2 is *Rhipicephalus sanguineus* according to sequence identity.

### DNA Extraction

Individual ticks were washed with phosphate-buffered saline, air dried for 10 min on tissue paper and separately sliced into small pieces by a sterile scalpel blade. They were then manually homogenized with a sterile micro-pestle, re-suspended in 200 μl of lysis buffer and 20 μl of proteinase K (20 µg/ml stock solution). After overnight incubation at 56°C with continuous gentle shaking, DNA was extracted using the QIAamp DNA tissue extraction kit (Qiagen, Hilden, Germany), following the manufacturer’s protocol. Eluted DNA (100 μl) was stored at −20°C until further use.

### 16S amplicon analyses

Forty tick-DNA pools were created, each containing equal concentrations of DNA from five ticks isolated from the same region, host and tick species. Tick pools were named after region and host and then given numerical identities (e.g. Nablus Dog Tick Pool 1, Nablus Dog Tick Pool 2). Tick pools were subjected to 16S rRNA sequence analyses on the Illumina platform, according to the manufacturer’s instructions [37]. In brief, V3–V4 hyper-variable regions were amplified using primers tagged with Illumina-specific primers. Amplification was performed using X2 KAPA HiFi HotStart ReadyMix (Kappa Biosystems). PCR conditions were as follows: 95°C for 3 minutes, followed by 25 cycles of: 95°C for 30 seconds, 55°C for 30 seconds, 72°C for 30 seconds, then 72°C for 5 minutes and final hold at 4°C. Negative controls containing nuclease-free water were used in each PCR run. PCR products were purified using AMPure XP beads (X0.8) followed by the second round of amplification that introduces the index sequences using the Nextera XT Index Kit (Illumina). Amplicons were sequenced according to the manufacturer’s recommendations. The PhiX control (15%) was added to the denatured library. Paired-end sequencing was carried out on the Illumina MiSeq platform.

16S rRNA amplicon sequences were analysed via the QIIME1 pipeline [38]. Amplicon sequences were demultiplexed into samples by barcode, then pre-processed and quality-filtered, using QIIME’s multiple_join_paired_ends.py and split_libraries_fastq.py scripts, accepting only those with a quality score below 19 and a minimum sequence length of 350 bp. Sequences passing the quality filter were clustered at 97% homology using QIIME’s pick_open_reference_otus.py and pick_otus.py scripts with the option “enable_rev_strand_match True”. Operational taxonomic units (OTUs) were defined at 97% identity and then taxonomically assigned using the Greengenes 13_8 database [39]. To remove spurious sequences, OTUs that occurred only once in the data were removed. We then calculated relative abundances for each OTU across the taxonomic levels from phylum to family, focusing on taxonomic sequence identifications relevant to tick-borne potential pathogens and endosymbionts (S1 Table).

### Shotgun metagenomic sequencing

Fourteen individual tick DNA samples were selected for follow-up by shotgun metagenomics, named after the pool they belonged to with a final numerical identifier for each tick, e.g. Nablus sheep tick 4.1, Nablus sheep tick 4.2. DNA sequencing libraries were prepared using the Nextera XT kit (Illumina), with DNA fragmented, tagged, cleaned and normalized according to the manufacturer’s recommendations. The quality of DNA in each library was evaluated using the Agilent 2200 TapeStation (Agilent) and the quantity was measured using the Qubit (Invitrogen). Libraries were sequenced on a NextSeq 500 with the mid-output setting and single-end reads. We analysed shotgun metagenome sequences using the Cloud Infrastructure for Microbial Bioinformatics [40]. Sequences were assessed for quality using FastQC version 0.11.5 [41] and quality-filtered using Trimmomatic version 0.35 with default parameters [42]. Trimmomatic’s Illuminaclip function was used to clip off Illumina adapters. Metagenome sequences were deposited in the Sequence Read Archive under Bioproject XYZ.

### Sequence-based identification of tick and host

Reads from each metagenome were assembled into contigs and then subjected to BLAST searches with mitochondrial signature genes from the potential tick and host species (S2 Table). Coding sequences from the metagenomic assemblies that had been identified from BLAST hits were then subjected to a second round of BLAST searches to determine the percentage identity to signature genes from relevant taxa.

### Taxonomic profiling of tick metagenomes

We performed a taxonomic assignment of reads in each sample using the Centrifuge metagenome classifier Version 1.0.3 together with its associated p+h+v database [43]. Centrifuge reports were converted to Kraken-style reports using Centrifuge’s kreport parameter and Pavian was used to visualize the reports [44]. Taxa with a relative abundance of <5% were discarded as potential kit contaminants and we then evaluated the reports for the presence of potential pathogens and endosymbionts.

### Assembly of microbial genomes

We constructed a set of completed reference genomes for each genus of potential pathogen or endosymbionts identified in at least one of our samples using Centrifuge (S3 Table). Reference sequences were downloaded using the ncbi-genome-download script [45]. We then mapped the metagenome from each sample against all the reference genomes using BowTie2 version 2.3.4.1 [46]. We converted SAM files to sorted BAM files using SAMtools and visualized stats using Qualimap 2 [47,48]. To allow us to bin species-specific reads for each microorganism of interest from the metagenomic sequences, the set of metagenomic reads from each sample that mapped to a taxon of interest were assembled into contigs using SPAdes (version 3.11.1) [49] and annotated using Prokka (version 1.12). We then used taxonomic profiles from Centrifuge for each contig to confirm the specificity of our binning approach. The coverage of the resulting draft genome sequences was calculated after mapping reads back to the assemblies using BowTie2. We compared the gene content between our metagenome-derived draft bacterial genomes and related reference genomes using Roary [50]. To confirm species identity, average nucleotide identity was calculated from BLAST searches [51] or by using the online ANI/AAI matrix tool [52].

### Phylogenomic analyses

We aligned signature genes from selected microbial taxa (*Supplementary files*) with homologues in metagenome-derived genome sequences using MAFFT (version 7.305b) [53] and performed maximum likelihood phylogenetic reconstructions using RAxML (version 8.2.12) with 1000 bootstraps [54]. MEGA7 was used to visualize the trees [55]. FastTree2.1 was used to construct phylogenetic trees using Generalized time reversible (GTR) and the CAT approximation (estimated rate of evolution for each site) model [56]. Bootstrapping with 1000 iterations was used to evaluate the significance of branches within phylogenetic trees. SNP distance matrices between assemblies and reference genomes were calculated using Snippy version 3.1 incorporating Freebayes version 1.1.0 for SNP detection [57,58].

## Results

### Taxonomic profiling of pooled tick samples

Two hundred hard ticks were collected from diverse animal hosts (23 dogs, 7 sheep, and 2 camels) and were identified morphologically to the species level (Fig. 2). PCR-amplification of 16S rRNA sequences was attempted on 40 pools of tick DNA. No products were obtained from eight of the pools. Sequencing of amplification products from the remaining 32 pools identified the major residents of the tick microbiome at the level of bacterial family (S1 Table). Four families harbouring potential pathogens or endosymbionts accounted for around 45% of all such sequences: the *Coxiellaceae, the Francisellaceae, the Rickettsiaceae* and the *Anaplasmataceae* (Fig. 3).

**Fig. 3.**
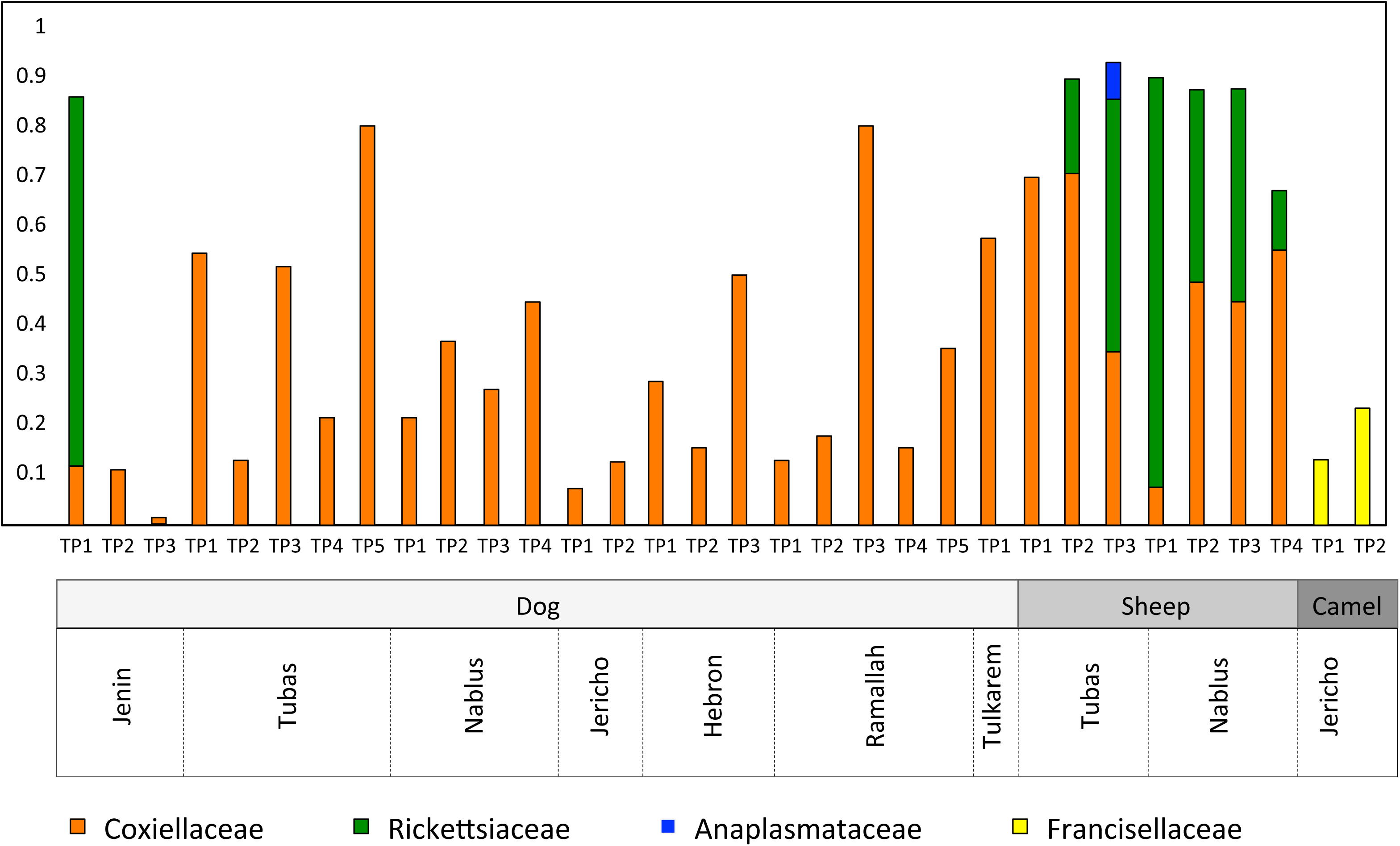
Relative abundances of potential pathogens and endosymbionts in pooled tick samples.

As expected from previous reports [33,34], the *Coxiellaceae* were detectable in all dog and sheep ticks, although their relative abundance varied from <10% to >80%. The *Rickettsiaceae* were detected as major residents (12–83% of reads) in all the sheep tick pools and in one tick pool derived from dogs. One sheep-derived tick pool also contained reads from the Anaplasmataceae. The two pools derived from ticks isolated from camels were, in line with previous work [12], dominated by the *Francisellaceae,* but lacked sequences from the *Coxiellaceae*.

### Metagenomic profiling of individual ticks

Fourteen ticks were selected for follow-up studies on the basis of the 16S results: two camel ticks, five sheep ticks, six dog ticks from the genus *Rhipicephalus* and two dog ticks from the genus *Haemaphysalis*. Shotgun metagenome sequencing of DNA from the fourteen ticks generated 15.8 million single-end reads, with >98% of reads from each sample surviving trimming (S5 Table).

Sequence-based identification of the ticks based on mitochondrial signature gene sequences generally confirmed the morphological identifications, with closest BLAST hits typically showing sequence identities of >99% to sequences assigned to the relevant tick species (Table 1). However, there were three discrepancies: Jericho Camel Tick 2.1 identified on morphological grounds as *Hyalomma impeltatum* was re-assigned on grounds of sequence identity to *Hyalomma dromedarii*; Nablus Sheep Tick 2.1 was identified morphologically as *Rhipicephalus turanicus,* but showed greatest sequence similarity to *Rhipicephalus sanguineus* dog ticks from Egypt and Iran; and Ramallah Dog Ticks 1.1 and 1.2, assigned to the species *Haemaphysalis parva* by morphology, both delivered a highest-scoring BLAST hit to a sequence from the species *Haemaphysalis concinna*, but the low level of sequence identity (88%) suggests that these ticks belong to a previously uncharacterised species within the genus *Haemaphysalis*. Identification of the tick host via mitochondrial signature gene sequences proved successful in six of the fourteen individual ticks (Table 1). This varied success presumably reflects the variation in the state of engorgement of the tick with the host’s blood.

**Table 1.** Metagenomic analyses of individual tick samples from Palestine.

| Tick | Tick ID by morphology | Tick ID by sequence (% identity to closest hit) | Host ID by sequence | Key microbes in taxonomic profile | Microbe ID from metagenome |
| --- | --- | --- | --- | --- | --- |
| Jericho Camel Tick 2.1 | *Hyalomma impeltatum* | *Hyalomma dromedarii* (100) | undetermined | *Francisella* | *Francisella persica* |
| Nablus Sheep Tick 1.1 | *R. turanicus* | *R. turanicus* (99.2) | undetermined | *Coxiella*, *Rickettsia* | *Rickettsia barbariae* strain Rb Nablus |
| Nablus Sheep Tick 2.1 | *R. turanicus* | *R. sanguineus* (99.1) | *Ovis aries* | *Coxiella* | undetermined |
| Nablus Sheep Tick 3.1 | *R. turanicus* | *R. turanicus* (99.2) | undetermined | *Coxiella*, *Rickettsia* | *Coxeilla mudrowiae* strain CRt Nablus<br>*Rickettsia massiliae* strain Rm Nablus |
| Tubas Sheep Tick 3.1 | *R. turanicus* | *R. turanicus* (99.3) | undetermined | *Coxiella*, *Anaplasma* | *Anaplasma ovis* |
| Tubas Sheep Tick 3.2 | *R. turanicus* | *R. turanicus* (99.2) | undetermined | *Coxiella*, *Anaplasma* | *Anaplasma ovis* |
| Hebron Dog Tick 8.1 | *R. sanguineus* | *R. sanguineus* (99.4) | *Canis lupus familiaris* | *Coxiella* | undetermined |
| Ramallah Dog Tick 1.1 | *Haemaphysalis parva* | *Haemaphysalis concinna* (87.7) | undetermined | *Coxiella* | undetermined |
| Ramallah Dog Tick 1.2 | *Haemaphysalis parva* | *Haemaphysalis concinna* (87.8) | *Canis lupus familiaris* | *Coxiella* | undetermined |
| Nablus Dog Tick 1.1 | *R. sanguineus* | *R. sanguineus* (99.4) | undetermined | *Coxiella* | undetermined |
| Nablus Dog Tick 1.2 | *R. sanguineus* | *R. sanguineus* (99.2) | *Canis lupus familiaris* | *Coxiella* | undetermined |
| Tubas Dog Tick 2.1 | *R. sanguineus* | *R. sanguineus* (99.2) | *Canis lupus familiaris* | *Coxiella*, *canine parvovirus* | Canine parvovirus |
| Tubas Dog Tick 3.1 | *R. sanguineus* | *R. sanguineus* (99.3) | undetermined | *Coxiella* | *Coxeilla mudrowiae* strain CRs Tubas |
| Tubas Dog Tick 3.2 | *R. sanguineus* | *R. sanguineus* (99.9) | *Canis lupus familiaris* | *Coxiella* | undetermined |

When subjected to taxonomic profiling, 12.6 % of the metagenomic reads were classified as bacterial and 0.015 % of reads as viral (S7 Table). The taxonomic profiles confirmed the presence of the bacterial genera *Coxiella*, *Francisella*, *Rickettsia* and *Anaplasma* in the relevant ticks (Fig. 4). In addition, reads from canine parvovirus were identified in Tubas Dog Tick 2.1.

**Fig. 4.**
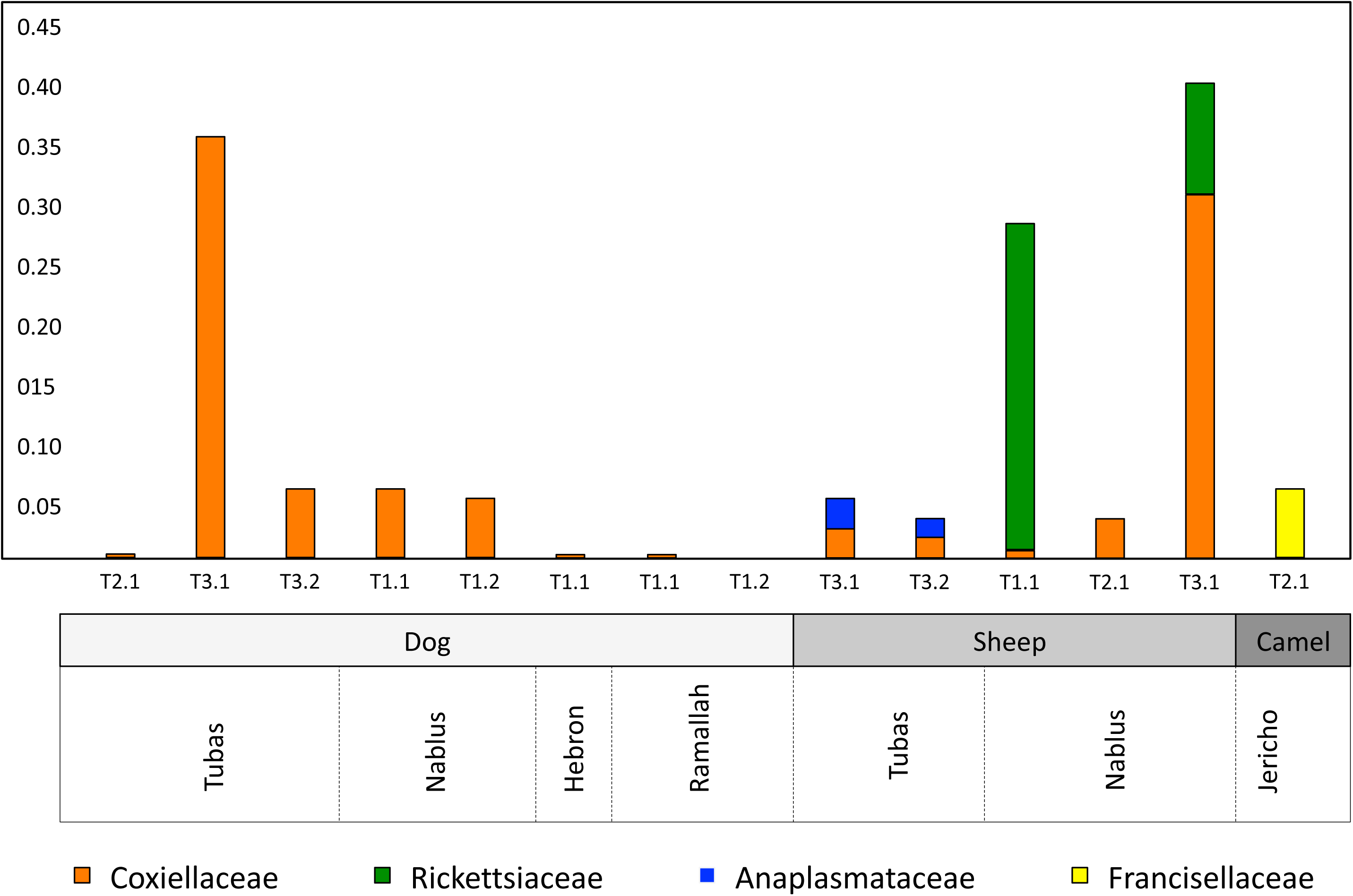
Relative abundance of potential pathogens and endosymbionts in individual tick samples

### Phylogenomic characterisation of tick-borne potential pathogens

Taxonomic profiling showed a high abundance of reads assigned to the genus *Rickettsia* in Nablus Sheep Tick 1.1 and Nablus Sheep Tick 3.1, which are both *R. turanicus*. In Nablus Sheep Tick 3.1, the *Rickettsia* reads (∼8%) were outnumbered by those from *Coxiella* (∼30%). However in Nablus Sheep Tick 1.1, the relative abundance of *Rickettsia* reads (∼25%) was much higher than for *Coxiella* (∼2%). Metagenomic reads from these two ticks were mapped against a collection of all complete rickettsial genome sequences (S3 Table). Reads mapping to any of the rickettsial genomes were extracted from each sample and were assembled into a draft genome sequence.

Rickettsial reads from Nablus Sheep Tick 3.1 assembled into 380 contigs, with an average depth of coverage of 13X, a genome size of 1,342,424 base pairs and a GC content of 32.73%. Comparisons to the *Rickettsia* reference genomes showed that this metagenome-derived genome (which we term “Rm Nablus”) was highly similar to the genome of *R. massiliae* strain MTU5 from Marseille, France [59], showing extensive synteny and just 127 SNPs spanning the whole genome (Fig. 5, S8 Table). The two strains also share a plasmid, *pRMA*. This high degree of genome conservation between *R. massiliae* strains from opposite ends of the Mediterranean region seems remarkable. However, both MTU5 and Rm Nablus strain are far less closely related (SNP distance >16000) to the only other genome-sequenced strain assigned to the species *R. massiliae*—the AZT80 strain from Arizona [60]. Instead, our comparisons show that this US isolate is much more closely related (just 4 SNPs different) to the strain ECT, which has been assigned to the species *R. rhipicephali*.

**Fig. 5.**
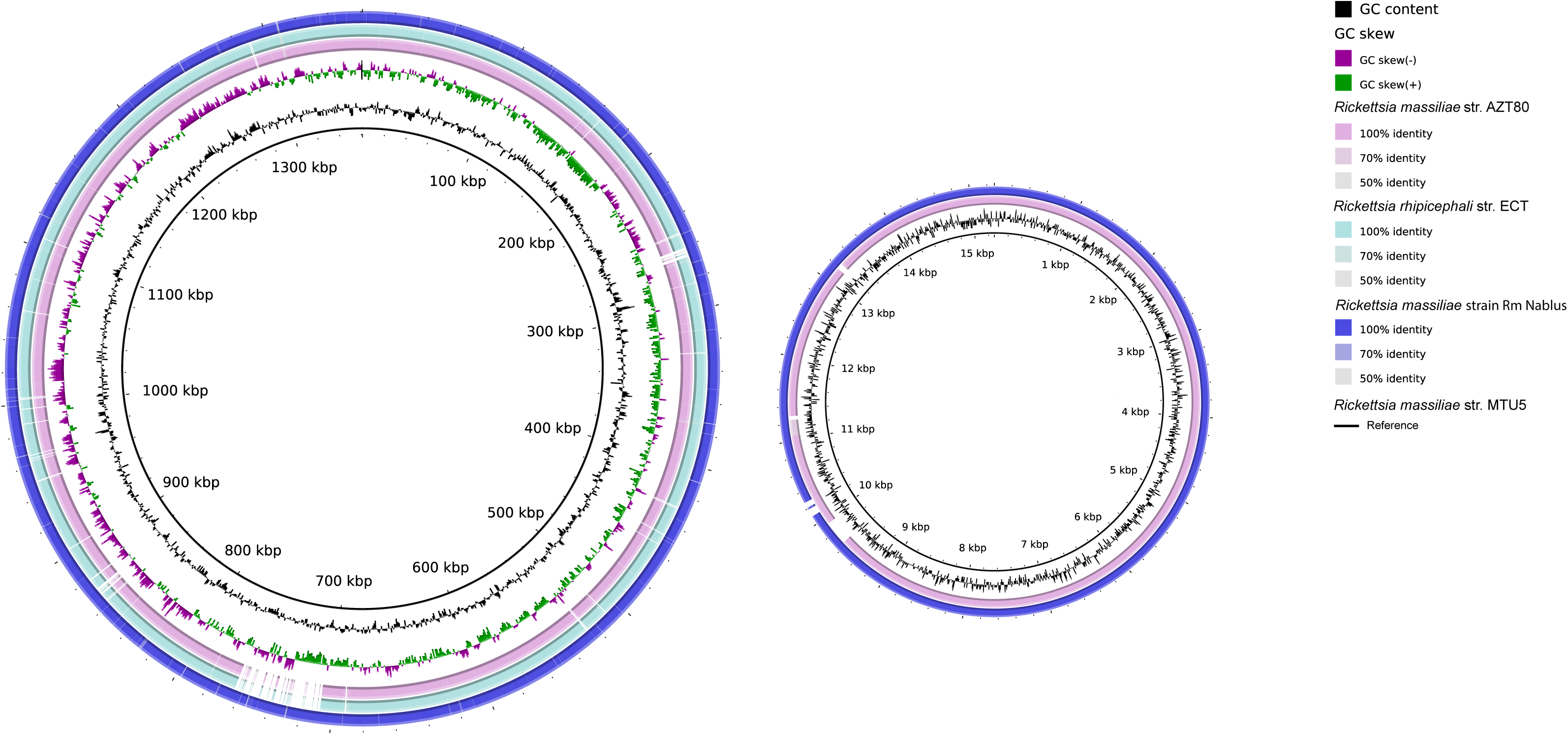
Genome comparisons for *Rickettsia massiliae* strain Rm Nablus. *From centre outwards: Rickettsia massiliae* strain MTU5 (black circle: GCF_000016625); GC content; GC skew; *Rickettsia rhipicephali* strain ECT (GCF_000964905), *Rickettsia massiliae* strain AZT80; *Rickettsia massiliae* strain Rm Nablus (this study).

Rickettsial reads from Nablus Sheep Tick 1.1 assembled into 47 contigs with an average depth of coverage of 14X, a genome size of 1,246,042 base pairs and GC content of 32.4%. Comparison with other rickettsial genome sequences showed that the closest genome-sequenced relatives of the *Rickettsia* from Tick 9 are *R. africae* str. ESF-5 and *R. parkeri* str. AT24. However, each are separated from this genome by >10,000 SNPs (S2 Figure), suggesting that it represents a distinct species for which no genome sequence yet exists. With that in mind, we performed comparisons to sequences from the unidentified genome to a database of rickettsial signature gene sequences that have been widely used to identify isolates to species level. Here, we found 100% identity with the 16S, *gltA* and *recA* genes from *Candidatus* R. barbariae and 582/584 residues identical in *ompB*. This leads us to conclude that our metagenome-derived assembly (which we term “Rb Nablus”) represents the first draft genome sequence from this species. Comparisons with the *R. africae* and *R. parkeri* genomes suggest extensive synteny between all three genomes. The only major exception appears to be the variable *tra* region, first identified in *R. massiliae* [59], where *Candidatus* R. barbariae lacks the *tra* genes found in the other two species (Fig. 6).

**Fig. 6:**
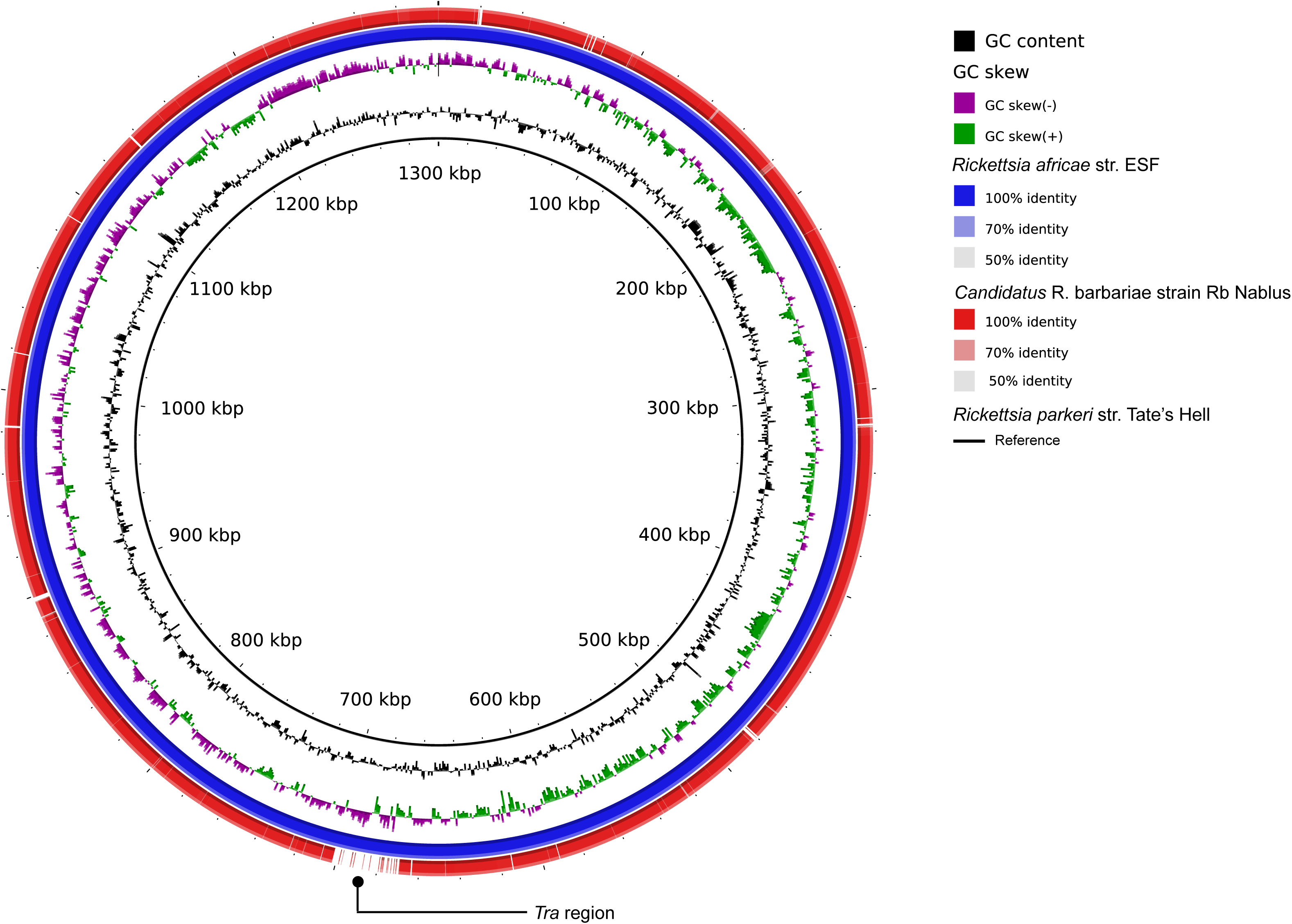
Genome comparisons for *Candidatus* Rickettsia barbariae strain Rb Nablus. *From centre outwards: Rickettsia parkeri* strain Tates Hell (black circle: NCBI ID GCF_000965145); GC content; GC skew; *Rickettsia africae* strain ESF-3 (GCF_000023005); *Candidatus* Rickettsia barbariae strain Rb Nablus (this study).

Taxonomic profiling indicated a high abundance of reads assigned to the genus *Anaplasma* in Tubas Sheep Ticks 1.1 and 1.2. Metagenomics reads from these two ticks were mapped to a set of 44 completed *Anaplasma* genomes (S3 table). Reads mapping to any *Anaplasma* genome were extracted and assembled into contigs. The *Anaplasma* assembly from Tubas Sheep Tick 1.1 contained 155 contigs with a total length of 312,716 bp, while that from Tubas Sheep Tick 1.2 contained 323 contigs with a total length of 497,831 bp. BLAST comparisons to the *Anaplasma* reference genomes showed that contigs from both ticks were highly similar (>99% sequence identity) to the genome of *Anaplasma ovis*.

As some reads from Tubas Dog Tick 2.1 were taxonomically assigned to canine parvovirus, we mapped this metagenome against the reference sequence for this virus (NCBI Reference Sequence: NC_001539.1) and obtained 11X coverage of canine parvovirus strain Tubas. BLAST comparisons with other canine parvovirus gene sequences showed that the Tubas Dog Tick 2.1 virus was most closely related to the lineage 2b strain CPV-LZ2 from Lanzhou in North-West China.

### Phylogenomic characterisation of tick endosymbionts

Taxonomic profiling showed a high abundance (>30%) of reads assigned to the genus *Coxiella* in Nablus Sheep Tick 3.1 and Tubas Dog Tick 3.1, so we mapped metagenomic reads from these ticks against a set of all completed *Coxiella* genomes (S3 Table). We retrieved all threads that mapped to any *Coxiella* genome from these samples and, as before, performed genome assemblies.

*De novo* assembly of *Coxiella* reads from Nablus Sheep Tick 3.1—a specimen of *R. turanicus*—resulted in 35 contigs with a combined size of 1,581,648 base pairs and a GC content of 38.2% and a depth of coverage of 37X. Phylogenetic analyses showed that the resulting draft genome (which we term “strain CRt Nablus”) was almost identical (with only 294 SNPs spanning the whole genome) to the Coxiella-like endosymbiont *Candidatus* Coxiella mudrowiae, strain CRt (ASM107771v1), derived from a *R. turanicus* sample collected at Kibbutz Hulda, Israel, 56 km from Nablus.

*De novo* assembly of Coxiella reads from Tubas Dog Tick 3.1 resulted in 177 contigs with a combined size of 1,529,439 base pairs and a GC content of 38.1%. The resulting draft genome (which we term “strain CRs Tubas”; Fig. 8) is relatively distantly related (>20,000 SNPs or an average nucleotide identity of only 98%) to the only other known genome of a *Coxiella*-like endosymbiont from *Rhipicephalus sanginueus*—strain CRs (NZ_CP024961.1)—which was derived from a sample collected in Caesarea, Israel, 48 km from Tubas. However, signature genes from our strain CRs Tubas all showed >99.5% nucleotide identity to those from other *Coxiella*-like endosymbionts from *Rhipicephalus turanicus*, while they showed only ≤96% identity to the same genes from similar endosymbionts derived from *Rhipicephalus sanguineus* (S. 10 Table)—suggesting that endosymbionts from the these tick species are vertically inherited and co-evolving with their hosts.

**Fig. 8.**
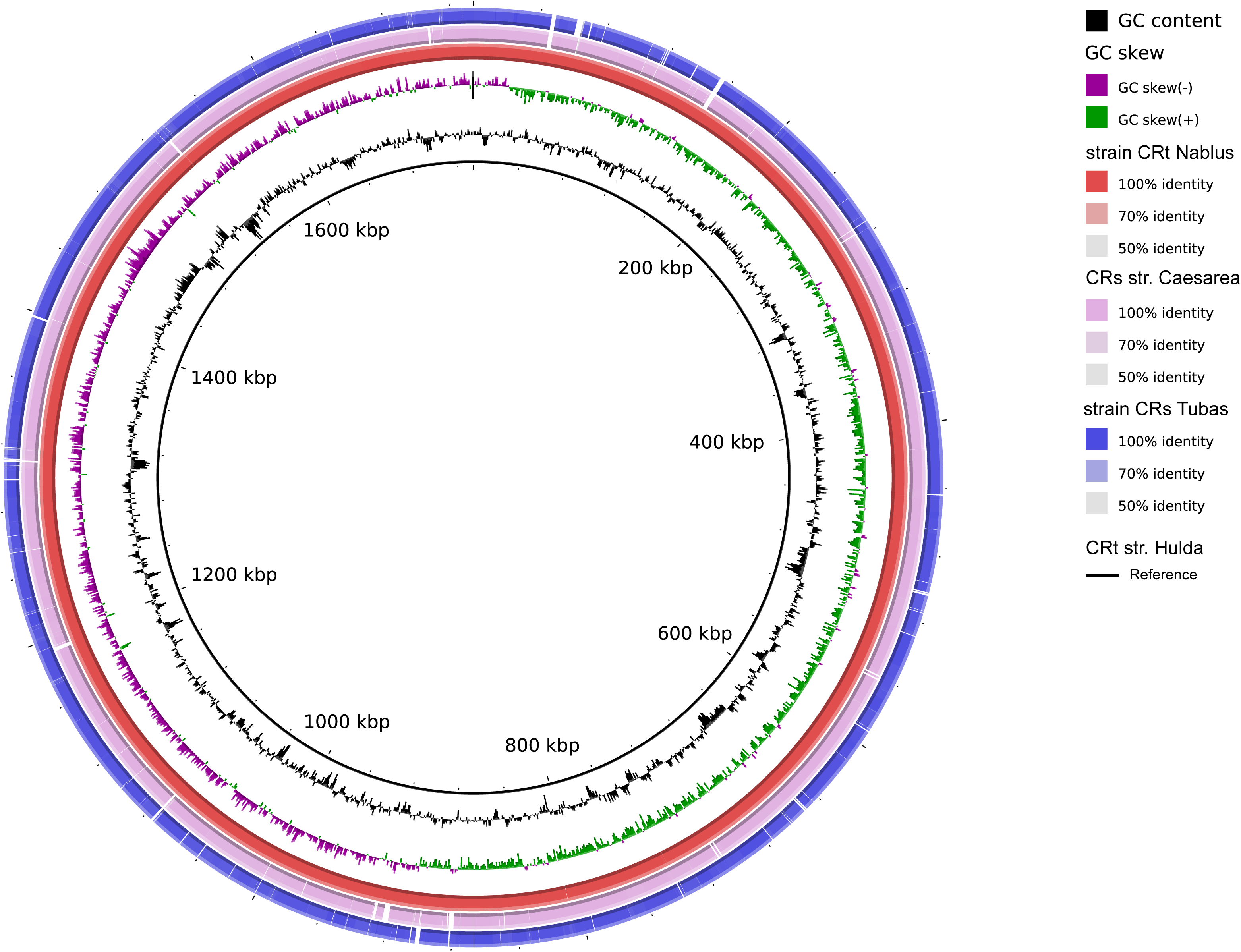
**Genome comparisons for** *Candidatus* Coxiella mudrowiae **strains CRt Nablus and CRs Tubas.** Fig. 8 Legend.

Taxonomic profiling of metagenomic reads from Jericho Camel Tick 2.1, a specimen of the species *Hyalomma dromedarii,* confirmed the presence of *Francisella*-like endosymbionts. However, when these reads were mapped against relevant *Francisella* genomes the depth of coverage was insufficient for further phylogenetic analysis.

## Discussion

Here, we have shown how a cost-effective combination of 16S rRNA sequence analysis and shotgun metagenomics can be used to detect and characterise tick-borne pathogens and endosymbionts. According to morphological criteria, we surveyed ticks from four different taxa [61]: the brown dog tick *Rhipicephalus sanguineus,* which has a global distribution [62]; its close relative *Rhipicephalus turanicus,* which we collected from sheep; a tick from the genus *Haemaphysalis,* collected from a dog; and *Hyalomma* samples from camels. All these taxa have been reported to feed on humans [61]. Interestingly, metagenomic sequencing allowed us to confirm or refine the species identities assigned to the ticks on morphological grounds, as well as in some cases confirming the identity of the host.

Mindful of the potential for reagent-based contamination of low-biomass samples [63], here we have focused on taxa already known to contain tick-borne pathogens and endosymbionts, rather than attempting to provide a definitive survey of tick microbiomes. Our metagenomics strategy has delivered three important benefits. First, in using sequence-based methods to confirm the identities of ticks (and their hosts), we avoid the problems of using morphological criteria alone, which can be problematic in tick identification and classification [61]. Second, in addition to simply detecting or identifying tick-borne pathogens and endosymbionts, we, like others [34,36], have used shotgun metagenomic sequencing to obtain in-depth genome-resolved characterisation of key bacteria from this setting. Third, in recovering genome sequences from an unexpected pathogens (canine parvovirus) and a previously unsequenced pathogen (*R. barbariae*), we have demonstrated the open-ended nature of diagnostic metagenomics [35].

*Rickettsia massiliae* has caused tick-borne spotted fever in humans in the Europe and South America [64–66]. Although no cases of human infection have been described in the Levant, this pathogen has been detected in ticks collected in Palestine and in Israel [24,28,29,31,32]. *Candidatus* R. barbariae belongs to the spotted-fever group of rickettsiae [67], but has yet to be associated with human infection. However, it has been found in human-parasitising ticks [68] and may well be pathogenic for humans. This candidate species has been detected previously in ticks from Palestine, from Israel and from Lebanon [24,29,32,69].

Here, we provide a third draft genome from *Rickettsia massiliae* and the first draft genome from *Candidatus* R. barbariae, which confirms its close similarity to the established human pathogens, *R. africae* and *R. parkeri* and provides a starting point for future population genomics studies of this potential pathogen. Comparison with pre-existing examples reveal that our *Rickettsia massiliae* Rm Nablus strain shows far greater sequence similarity to the MTU5 strain from Marseille than to the AZT50 strain from Arizona, hinting at an Old World/New World phylogeographic population structure for this pathogen, which awaits clarification from additional genome sequences.

Coxiella-like endosymbionts of *Rhipicephalus sanguineus* and *R. turanicus* have been recently assigned to a new candidate species *Candidatus* C. mudrowiae, on the basis of genome sequences derived from pools of ticks isolated in Israel [33,34]. We present genome sequences of *Candidatus* C. mudrowiae derived from individual *Rhipicephalus sanguineus* and *R. turanicus* ticks, which were harvested in Palestine. Unsurprisingly, *Candidatus* C. mudrowiae genomes from Palestine and Israel are very similar. Interestingly, our two genome assemblies are each clearly more closely related to one of the variants in each of the tick pools from Israel, pointing the way towards a more sophisticated genome-informed documentation of phylogeographic variation in this species, which will be relevant to the analysis of host and endosymbiont co-evolution.

We detected two microorganisms by metagenomics that were somewhat unexpected: *Anaplasma ovis* and canine parvovirus. *Anaplasma ovis* infects sheep and goats in many regions of the world, but its precise prevalence and clinical importance remains uncertain in most settings, including here in Palestine [70]. Enteritis caused by canine parvovirus has been a leading cause of morbidity and mortality in dogs globally since it emerged as a new pathogen in the mid-1970s [71]. The usual route of transmission is thought to be faecal-oral. However, the infection is characterised by a marked viraemia and canine parvovirus has been shown to persist within hard ticks under laboratory conditions [72]. Our metagenomics survey has provided the first evidence that ticks can carry the virus in the natural setting and raises the possibility that ticks might contribute to the spread of canine parvovirus among dogs in Palestine and more widely. However, it remains possible that the tick is merely a dead-end host for this virus.

In closing, the results presented here hint at exciting opportunities for more ambitious metagenomic surveys in the future, as sequencing technologies become more user-friendly and cost-effective. Such ventures are likely to shed new light on the transmission, evolution, and phylogeography of tick-borne pathogens and endosymbionts.

## Supporting information

S1 Table: 16S OTU table for tick pools

S2 Table: Tick and host mitochondrial sequences

S3 Table: Reference genomes for tick-borne pathogens and endosymbionts S4 Table: Results of targeted PCRs on individual ticks

S5 Table: Shotgun Sequencing Statistics

S6 Table: Mapping to tick and host mitochondrial sequences

S7 Table: Taxonomic profiles of metagenomic sequences

S8 Table Mapping data for tick-borne pathogens and endosymbionts

S9 Table: SNP Tables for tick-borne pathogens and endosymbionts

S1 Figure. Multiple alignment of signature sequences confirming identity of *Candidatus* R. barbariae Rb Nablus

### Acknowledgments

We thank Yuval Gottlieb for reading and commenting on the manuscript.

## Author Contributions

Conceived and designed the experiments: SE, AN, and MP. Identified the ticks and extracted DNA: AA-J and OA-S. Performed 16S PCR amplification: SE and AN. Performed 16S amplicon sequencing: OA-S and HH. Performed metagenomic sequencing: SE and AN. Bioinformatics analysis: AR and MP. Wrote the manuscript SE, MP, AR and AN. All authors approved the manuscript.

